# Fronto-Hippocampal Synchronization in Rapid Spatial Learning in Humans

**DOI:** 10.1101/2025.11.07.685935

**Authors:** Maya Geva-Sagiv, Soyeon Jun, Marta Kryven, Josh Tenenbaum, Erie D. Boorman, Randy O’reilly, Jack J. Lin, Charan Ranganath, Ignacio Saez

## Abstract

Achieving goals in real-life situations—from fetching a glass of water to landing a dream job—often requires planning based on experience and executing a sequence of actions. Neurophysiological research in animal models has indicated that the orbitofrontal cortex (OFC) mediates relationships between memory, actions, and outcomes and the hippocampus and other medial temporal lobe (MTL) regions are known to be critical for rapid learning, but little is known about how these areas interact to support rapid learning and retrieval of goal-directed action sequences in humans. Here, we leverage a rare opportunity to investigate human OFC gamma oscillations and examine the coordination between the OFC and MTL during a continuous multi-step task that requires applying recently acquired experience to guide behavior. We used multisite intracranial electroencephalography (iEEG) recordings while participants searched for a hidden goal in an animated game to study neural activity in both brain areas during goal-directed behavior. Hippocampal ripples—brief high-frequency oscillations reflecting synchronized neuronal firing—are known to support memory consolidation during sleep, but their role during active memory retrieval and updating remains unclear. We found that OFC gamma activity was modulated by both memory demands and ripples in the hippocampus and adjacent structures. Notably, ripple-coupled OFC gamma during exploration was associated with subsequent task performance. We propose that hippocampal ripples mark a narrow window, supporting hippocampal-cortical communication required for successful goal encoding for future behaviors.

## 1 Introduction

Many real-world tasks, such as planning and navigation, involve a series of computations combining prior knowledge and real-time experience of a constantly changing environment. In this work, we focus on bi-directional interactions between prefrontal regions supporting cognitive control and the medial temporal regions supporting declarative memory, hypothesized to be critical for multi-step decision making (Simons & Spiers, 2003). Theories of hippocampal activity that focus on spatial cognition often imply that the hippocampus is constantly encoding and retrieving spatial information to convey to the cortex (Byrne et al., 2007). In contrast, functional magnetic resonance imaging (fMRI) research on human episodic memory suggests that bi-directional hippocampal-prefrontal interactions should primarily occur during computationally significant moments, such as event boundaries (Baldassano et al., 2017; Barnett et al., 2023; Franklin et al., 2020; Zacks, 2020, p. 200). It is therefore possible that, during multi-step goal-directed behavior, cortico-hippocampal interactions might be optimized to support memory processes associated with planning and goals.

Prior work with animal models suggest that hippocampal ripples—brief high-frequency oscillations—create selective windows for hippocampo-cortical information transfer by facilitating reinstatement of newly acquired information (Eliav et al., 2025; Foster, 2017; Tang et al., 2017; Todorova & Zugaro, 2020). Human studies show ripples precede successful memory retrieval and correlate with neocortical activity during recollection retrievals (Henin et al., 2021; Norman et al., 2021; Vaz et al., 2020), but the role of hippocampal-cortical interactions across distinct behavioral phases -- including exploration, planning, execution and goal attainment remains little explored.

The orbitofrontal cortex (OFC) is critical for contingent learning—recognizing stimulus-outcome relationships and updating values when goals are reached (Boorman et al., 2021; M. P. H. Gardner & Schoenbaum, 2021; Rudebeck & Murray, 2014; Rushworth et al., 2011; Saez et al., 2018; Walton et al., 2010). The hippocampus, which shares reciprocal connectivity with OFC (Barbas & Blatt, 1995; Swanson et al., 1978; Wikenheiser & Schoenbaum, 2016), reinstates neural patterns associated with both spatial paths and non-spatial associative relationships (Koster et al., 2018; Park et al., 2020; Zeithamova et al., 2012). This hippocampal reinstatement may facilitate OFC coding of outcome-predictive behaviors (Biderman et al., 2020; Shohamy et al., 2009; Witkowski et al., 2025). In this framework, the hippocampus processes contextually-related events, cues, and outcomes, then sends this context-specific information to prefrontal cortex, where neural ensembles develop contextual rules during learning (Eichenbaum, 2017).

In this work, we examine the role of hippocampal-cortical interactions in a continuous spatial navigation and cognitive map learning task. We focus on two possible ways in which these interactions might occur. One possibility is that hippocampal-cortical interaction could, in theory, be continuous and sustained, for instance, through OFC continuously integrating a stream of hippocampal information for ongoing decision-making. For instance, theories of predictive cognitive maps implemented through updates to the successor representation (SR) -- the predictive maps encoding likely future states -- support this prediction because the SR provides a continuously updated estimate of future outcomes (Stachenfeld et al., 2017). Predictive cognitive maps have been documented on a behavioral level as well, with the predictive computations explained by a continuous process driven by Bayesian inference (Sharma et al., 2022).

Another possibility is that cortico-hippocampal interactions might be concentrated during particular phases, rather than continuously uniform. Consistent with this possibility, animal studies have demonstrated hippocampal-prefrontal synchronization was stronger during awake ripples and enhanced firing early learning (Tang et al., 2017). Human studies using maze navigation tasks, hippocampal activity peaked during planning epochs when participants had to consider sequential choices and predict future steps - i.e. at maze onset (Kaplan et al., 2017, p. 201). Further evidence for phasic computation peaks comes from a study where people navigated a virtual subway network, where both behavior and neural data were in line with a view that plans are formed and executed in a hierarchical fashion (Balaguer et al., 2016).

Testing competing models of hippocampal-prefrontal interactions—continuous spatial information processing versus selective engagement during planning and goal attainment— requires tasks that separate goal-selection from goal-engaged phases. Therefore, we use a continuous spatial navigation task, allowing us to identify distinct behavioral phases (exploration, planning, execution, goal attainment) and align neural activity with computationally significant moments. This approach is critical for determining whether hippocampal-prefrontal communication occur continuously or selectively during times that are useful for chunking continuous events into event segments (Baldassano et al., 2017; Ezzyat & Davachi, 2011; Gershman et al., 2014; Kurby & Zacks, 2008; Zacks, 2020). Using simultaneous intracranial EEG (iEEG) recordings from orbitofrontal cortex and medial temporal lobe in humans (Figure 1A), we investigated dynamics of OFC gamma oscillations and their modulation by different behavioral events and hippocampal (and adjoining area) ripples, while participants were performing an animated goal-seeking task (Kryven et al., 2024). We examined how these neural signatures evolve from exploration to experience-based behavior and whether they are functionally related to successful task learning.

**Figure 1.**
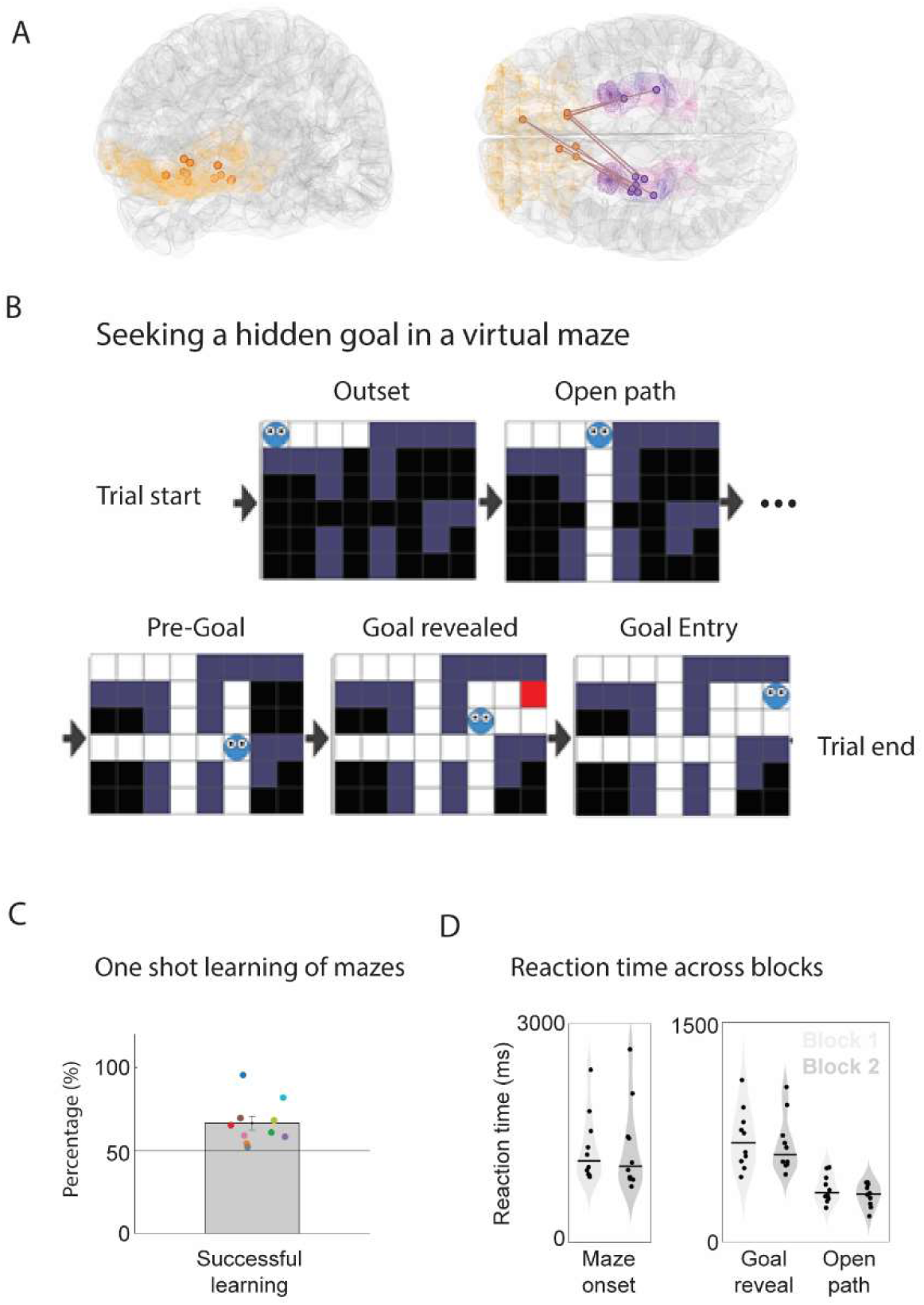
One-shot learning of hidden-goal locations in a virtual 2D Maze. **(A)** Participants are patients with simultaneous recordings from OFC (orange) and MTL regions (hippocampus, purple and amygdala, and parahippocampal gyrus, pink). Left: iEEG OFC contact locations are overlaid on a standard (Montreal Neurological Institute (MNI) brain template (see Methods for electrode localization). Right: Lines connect MTL-OFC pairs (both contacts from the same patient) included in coupling analysis (Figure 3). (**B**) Participants guided a virtual agent through a series of mazes to find the maze exit as efficiently as possible. Each maze featured a unique goal (exit) location (red square) and a spatial layout constrained by walls (gray squares). The goal location remained hidden (black squares) until the agent reached a position with a direct line of sight to that location (as shown in the ‘Goal revealed’ panel). Participants completed 30 different mazes per block, in randomized order across consecutive blocks. (**C**) The ratio of successfully learned mazes (over two blocks) was significantly higher than chance (n = 10 participants, Wilcoxon test - P = 0.002). Each dot depicts the percentage of successful paths per participant (also see Methods and Figure S2A). (**D**) Reaction time varied at different navigational events: patients’ response time was longer at complex navigational events, compared to simple moves in Open Path, likely reflecting deliberation before movement. Data points represent average reaction times (RTs) from individual participants. Horizontal lines mark group means for each behavioral-event type during block 1 (light gray) and block 2 (dark gray). Compared to moving from one square to the next in an open path (‘Open Path’), people took longer to move the agent when starting to navigate a new maze (‘Outset’), as well as when the goal was revealed (Goal Revealed). RT slightly decreased with learning, but differences were non-significant (Methods, note that only successfully learned trials are included in this analysis).

## 2 MATERIALS AND METHODS

### 2.1 Participants

Intracranial recordings were obtained from 10 patients with medically intractable epilepsy (2 females) while they were undergoing the pre-surgical iEEG evaluation with stereotactically implanted electrodes. The electrode placement was determined solely based on the individual patient’s clinical needs, and we recruited patients who had at least one electrode implanted in the hippocampus. The age of the patients ranged from 18 to 51 (mean = 31.3, SD = 10.5; Table S1). Data were collected at UC Davis Medical Center (Sacramento, CA) and UC Irvine Medical Center (Orange, CA). All experimental procedures were approved by the Institutional Review Board at each hospital and were fully described to the patients before they consented to participate in the study. Inclusion criteria for every analysis is detailed below based on location of electrodes for each patient and electrophysiological analysis – behavior (10 sessions), OFC gamma-power analysis (11 channels from 10 patients), and OFC-MTL coupling (11 couples from 6 patients).

### 2.2 Experimental design

Intracranial EEG signals were recorded while patients performed the Maze Search Task (MST), described in (Kryven et al., 2024). In the task, patients controlled a virtual agent to navigate a series of partially observable, two-dimensional mazes to locate a hidden exit (goal location) (Figure 1B). Every trial began with the onset of a maze on a screen, and the agent’s initial location was always the northwest corner of the maze. The player in the MST game could move the agent to any adjacent grid tile that is not blocked by walls (walls are marked as dark blue tiles, Figure 1B) in the four cardinal directions (up, down, left, right). The contents of the tiles paving the paths were initially occluded and became revealed part by part as the agent approached the unrevealed section. The exit (goal-location) becomes visible and marked as a red tile once the agent is close to it (goal-reveal location). The goal location varied across mazes: This feature of the maze kept the goal location unknown until the agent was near the goal location. Participants were told that each of the black tiles is equally likely to hide the exit. While trials were self-paced, participants were instructed to do their best to reach the exit as efficiently as they could. The trial ended when the participant moved the agent to the red-tile, which was 1-3 steps away from the goal-reveal location (see example in Figure 1B). Once the exit was found, the next maze was introduced. A block comprised of a series of 30 trials with unique mazes. Patients played 2-3 blocks, each comprising the same 30 unique mazes, which were presented once per block (Figure 1B) in randomized order. The mazes were set in grids of varying dimensions (e.g., 4x6, 8x7). Participants navigated mazes equally well (38%) or better (29%) in the second block, as assessed by the number of moves to reach the exit (Figure 1C). We defined successfully-learned mazes as those in which participants either reduced the number of steps or kept the same path as the first trial, and degraded mazes as those that had a longer path to exit on the 2^nd^ trial relative to the 1^st^ trial (see participant-specific breakdown in Figure S2A). This definition does not imply the path to the exit is the optimal one, only that the participant either retraced the original path taken or was able to improve it on the 2^nd^ try.

Participants used a keyboard, prompted on a table over the hospital bed, to move the agent on the screen, key strokes were recorded via Psychtoolbox (Matlab Mathworks) and were used to determine the timestamps for each behavioral event used for electrophysiological analysis (Table S2).

For analysis purposes - we split all locations on the maze according to their behavioral context, aggregating all locations that belong to the same behavioral-event regardless of their spatial location. See Table S2 for breakdown of the spatio-temporal characteristics of every behavioral event.

### 2.3 Behavior – Reaction time (RT) analysis

RTs at navigational locations, were compared against RTs of open-path moves, which we selected as a baseline for initiating a move in the maze for two reasons: 1) open-path moves did not involve a bifurcation or path/goal revelation; 2) Identified as the minimal RT in average when comparing different behavioral events on the maze (Figure S2B). RTs of other behavioral events (Maze Outset, Choice Point, Goal Revealed, and Goal Entering) were compared to RTs from neutral moves in separate mixed-effect regression models that included event type and block as predictors. The block variable was included in the models to account for potential response facilitation that correlates with time on task, using Matlab custom code:

~~~
model = fitlme(Data, ’RT ∼ ttype*block + (1|subj)’);
~~~

~~~
[beta, betanames, stats] = fixedEffects(model));
~~~

Bonferroni correction was used to correct for multiple comparisons (p = .05/4). RTs were z-scored within subjects before being entered into mixed-effect regression models.

### 2.4 iEEG recording and preprocessing

iEEG data were recorded using a multichannel research system (Neuralynx, https://neuralynx.com/) or a clinical monitoring system (Nihon Kohden, https://www.nihonkohden.com/). iEEG signals were acquired at a minimum sampling rate of 512 Hz while being referenced (online) to a subgaleal strip. Onsets of experimental events were recorded along with iEEG data via a photodiode placed at the bottom-left corner of the stimulus presentation screen.

iEEG data were preprocessed and analyzed using EEGLAB (Delorme & Makeig, 2004), Fieldtrip (Oostenveld et al., 2011), and custom MATLAB scripts. Data were offline re-referenced to the common average of signals from electrodes that met the following criteria: (1) placed in white matter, (2) mean amplitude of the electrode fell within 1.5 standard deviations of the mean signal across all channels, (3) absence of inter-ictal spikes, and (4) absence of prominent ERPs time-locked to trial events. The last criterion was adopted because re-referencing by common average signal, including prominent ERPs (potentially with conditional differences), can falsely introduce ERPs in all electrodes. Re-referenced signals were down-sampled to 512 Hz, notch-filtered for 60 Hz line noise and its 2^nd^, 3^rd^, and 4^th^ harmonics. Electrodes and epochs containing epileptiform activity were excluded after visual examination.

### 2.5 Electrode localization

Electrodes were localized to specific anatomical structures by aligning individual patients’ post-implant CT scans to the individual’s structural MRI scans, using Matlab LeGUI (Davis et al., 2021). Electrode locations were visually identified and marked in the individual’s native brain space, and the electrodes determined to be inside the hippocampal proper or in the orbitofrontal cortex (OFC) were included in the following iEEG analyses. The anatomical location of each electrode was determined using the Yale Brain Atlas (McGrath et al., 2022), which defines OFC subregions based on macroanatomical landmarks visible on MRI. These anatomical definitions do not correspond directly to cytoarchitectonic divisions such as Brodmann areas. For example, OFC 2B and 3B in the Yale atlas may roughly correspond to Brodmann area 11, while OFC 5B lies more ventrolaterally and may overlap with area 13. Given the known discrepancies between parcellation schemes, such correspondences are inherently approximate. While not directly comparing atlas systems, prior studies have highlighted the limitations of relying solely on cytoarchitectonic or macroanatomical divisions for defining cortical regions, particularly in the prefrontal cortex (Petrides & Pandya, 2007; Van Essen et al., 2012). A total of 18 electrodes were determined to be in the hippocampus, and 11 in the OFC (Figure S1).

### 2.6 Detection of ripple events

High-frequency ripples were detected from iEEG recordings from hippocampus, amygdala, and parahippocampal cortex using a modified version of previously published methods (Geva-Sagiv et al., 2023; Liu et al., 2022; Norman et al., 2019). All electrodes were bipolar referenced to a nearby white matter contact to eliminate common noise. Briefly, LFP signals from the hippocampal electrodes were filtered for the ripple band (70-180 Hz), and the instantaneous analytic amplitude was computed using a Hilbert transform. The instantaneous power amplitude of the resulting analytic signal was smoothed and converted into z-scores, and events, where the z-score exceeded 4 (i.e., 4 standard deviations (SD) above the mean), were detected as ripple events. Ripple events were considered to extend until the z-score fell under 2. Adjacent events that were within 30 ms of each other were merged into one event. Figure 3A depicts a grand average of ripples detected while performing the cognitive task as well as their spectral profile. The following events were excluded and not treated as SWRs: events that were shorter than 20ms or longer than 200ms, events that occurred within 50ms from an interictal epileptic discharge (IED; see below for IED detection), and events that coincided with increases (4 SD above the mean) of ripple-band power in the common average signals from all iEEG channels.

To distinguish between physiological ripples and other high-frequency oscillations (HFOs) related to epilepsy (Bragin et al., 2010), we discarded any candidate ripple event that occurred within 500 ms of inter-ictal epileptic discharges (IEDs). The detection of IEDs involved two complementary methods: an automated line-length algorithm based on (Baud et al., 2018) and a band-limited power analysis similar to (Smith et al., 2022). For the latter, the raw hippocampal iEEG signal was first filtered between 20-40 Hz using a zero-lag linear-phase Hamming windowed FIR filter. Next, the filtered signal underwent rectification, squaring, smoothing, and z-scoring. Events that exceeded 8 standard deviations (SD) were identified as IEDs.

After rejection of 7 channels with excessive pathological activity (from subjects 1,4-6, Methods), we included 9 channels (6 patients) from the hippocampus and surrounding areas (plotted in Figure 1A). We recorded n = 12812 ripples over 6 sessions of cognitive testing.

In order to assess how the rate of ripple occurrences changes around these empirical events, we constructed peristimulus time histograms (PSTH) of ripple occurrences that are time-locked to the events of interest (Figure S4A), similar to (Norman et al., 2019). PSTH were obtained for all 50-ms time bins that span the event time window, and the resulting bin-wise PSTH were smoothed by a 3-point triangular window (i.e., smoothed with 150-ms triangular window). Events that were associated with RT that fell outside 3 standard deviations from the mean in individual patient’s data were excluded from analyses. To test whether there were statistically meaningful fluctuations in the grand average trace from a given event (Figure S4A), we compared event-related PSTHs to surrogate data that were created by circularly jittering PSTH traces by random amount in each trial. For statistical testing on the ripple rate PSTH traces, a linear mixed-effects model was estimated for each time bin, and nonparametric cluster-based permutation tests were used to correct for multiple comparisons (Oostenveld et al., 2011). Regression estimates were compared against estimates that were obtained from conducting the identical models on permutated data. For testing the difference in ripple rates between blocks, block labels were shuffled for each permutation iteration. Final statistical testing used the 95^th^ percentile of the permutation distribution and cluster size as thresholds. This nonparametric cluster-based permutation test did not reveal significant increases in ripple rates in the behavioral events tested (maze onset, goal reveal, open path, pre-goal, or goal-entry).

To test modulation of OFC gamma-power around ripples, peri-ripple and non-ripple segments were matched by event-type and block, to establish OFC gamma increases are time-locked to ripples rather than general task timing. These analyses do not establish directionality; unmeasured arousal or movement states that may covary with ripple occurrence.

### 2.7 Analytic signals of oscillatory activity

The preprocessed LFP signals from the hippocampus and the OFC were spectrally decomposed as follows: continuous signals were filtered using FIR filters with frequency centers varying from 2 to 249 Hz. Frequency centers were spaced by bandwidth f ± 0.05 f from the adjacent frequency center (Ward & Doesburg, 2009), and this approach yielded 49 frequency centers. Filtered signals were Hilbert-transformed, and the instantaneous power amplitude and phase were computed from the resulting analytic signals.

### 2.8 Analysis of Oscillatory Activities in the OFC

#### 2.8.1 Gamma oscillations

Gamma oscillations (>30Hz) reflect synchronized neural circuit activity, facilitating inter-regional communication during decision-making and reward processing (Rich & Wallis, 2017; Siegel et al., 2012). While extensively studied in working memory tasks, their role during tasks involving longer periods of memory retention, such as multi-step planning tasks, remains unclear. The gamma frequency range of 50-100Hz used in our analysis was selected based on both established literature conventions and data-driven considerations specific to our dataset. Classical definitions of gamma oscillations vary across studies, with classical definitions related to general computations (Buzsáki & Wang, 2012a) defining the range as 30-90 Hz and intracranial EEG studies examining memory processes, such as Manning et al. (Manning et al., 2011), extending this to 30-99 Hz. Hippocampal-focused research typically employs lower frequency bounds to avoid contamination from high-frequency ripple oscillations (Colgin, 2015), while recent cortical gamma investigations have increasingly incorporated higher frequency components (up to 150Hz) (Kucewicz et al., 2018; Zhang et al., 2015). Time-frequency analysis of our data (Figure 2A) revealed that most spectral energy was concentrated below 100 Hz, supporting the inclusion of this frequency window. Importantly, our data-driven approach identified a significant cluster in the 32-102 Hz band during orbitofrontal cortex spectral analysis time-locked to ripple events, providing empirical justification for this specific range in our experimental context (Figure 3B). To ensure robustness of our findings, we validated our results using an extended frequency band (50-150 Hz), which yielded statistically similar outcomes, confirming that our chosen gamma range appropriately captured the relevant oscillatory dynamics without compromising the integrity of our analyses.

#### 2.8.2 Time-Frequency Power Analyses

Time-frequency response (TFR) power analysis was performed for aggregated epochs, triggered by reaching a specific behavioral timepoint (maze-onset, open-path, goal reveal, etc., see Figure 1B). TFRs (Figure 2A) were extracted by calculating a spectrogram around behavioral time points [-0.75,0.75]sec and subtracting it from the baseline (1sec) spectrogram. Spectrograms were calculated using *ft_specest_mtmconvol(),* (FieldTrip toolbox (Oostenveld et al., 2011), MATLAB, MathWorks, frequencies 1–250 Hz, 1-Hz resolution) using a sliding Hanning-tapered window with a variable, frequency-dependent length that comprised at least five cycles (Geva-Sagiv et al., 2023; Staresina et al., 2015). Time-locked TFRs of every behavioral event were then normalized as the percentage change from pre-event baseline and were averaged for each session. Averaged TFRs were then re-averaged, using a weighted average according to the number of trials for each behavior point in each session. To quantify changes in gamma-power, we defined a marker for detecting gamma-power increases time locked to different behavioral conditions ‘gamma-change percentage’ (GCP) - we calculated the average of TFR values in the gamma-range window (50-100Hz) during a [0.15-0.35] sec time-window locked to each behavioral timepoint.

We conducted statistical analysis to dissociate the effect of experience (blk) and behavioral events (event) by fitting a linear mixed effect model with GCP as a modeled factor as a function of behavioral event type, block and their interaction with random intercepts for subject (session). Statistical comparisons were conducted in MATLAB R2023 (using LinearMixedModel class, *fitlme*()) in the following manner:

~~~
formula = ’GCP ∼ event + blk + event*blk + (1|subj)’;
~~~

~~~
lme1 = fitlme(Data,formula);
~~~

~~~
anova(lme1);
~~~

~~~
betaTable = lme1.Coefficients;
~~~

Cohen’s d (Cousineau & Goulet-Pelletier, 2021), Figure S3) was calculated using *meanEffectSize*(), and plots were generated with *gardnerAltmanPlot*(), both from MATLAB R2023 (https://www.mathworks.com).

#### 2.8.3 Calculating lags using the amplitude cross-correlation method

We’ve used cross-correlation to detect changes in the temporal dynamics of response to behavioral locations between blocks 1 and 2 (Figure 2D (Adhikari et al., 2010)).

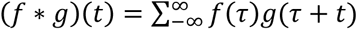

We generated a time course for each block/event type by averaging TFR during the studied window [0.15, 0.35] sec (for each block, see waveforms plotted in Figure 2A). We then calculated the correlation between these two time courses of gamma-power changes over lags ranging from -0.2 to 0.2 sec. The mean amplitude was first subtracted from each vector before cross-correlating them (Adhikari et al., 2010). The lag at which the cross-correlation peaked was then determined (xcorr()). A significant peak indicates there is a high correlation between the two time courses (indicated by the reported P value, using *corrcoef*()), and a shift of the center peak indicates there is a temporal shift between the two time courses (Adhikari et al., 2010). The significance of each block 1-block-2 gamma-band change cross-correlation was also verified using a shuffling procedure. For each event-type, time-courses were randomly permuted 1000 times. The permuted amplitude envelopes were then cross-correlated, yielding a distribution of cross-correlation peaks expected by chance. The original cross-correlation was considered significant if its peak value was greater than 95% of these randomly generated cross-correlation peaks.

### 2.9 Coupling of Oscillatory Activity in the OFC to Ripples

We conducted analyses to characterize the mid-prefrontal oscillatory activities that arise when ripples occur (see “Detection of ripple events” above). From each subject’s data, all possible pairs of ripple-contacts and OFC contacts were pooled and used for this analysis. (n = 11 electrode pairs from 6 sessions, Figure 1A, left). Instantaneous power amplitude data from the OFC were epoched for peri-ripple and non-ripple segments. Peri-ripple epochs were defined as - 200 ms to 200 ms from the peaks of hippocampal ripples, and non-ripple epochs were defined as 401-ms-long segments during which the hippocampal signal was free of ripples. To match the peri-ripple and non-ripple signals as closely as possible, the non-ripple epochs were matched with peri-ripple epochs in terms of the block and event type. Segments that included any IED were excluded from analyses. This procedure yielded a frequency x time x observation matrix of the instantaneous power amplitude for each condition (peri-ripple, non-ripple) and electrode-pair. The peri-ripple vs. non-ripple difference was estimated as paired-sample t-statistic for each frequency and time point. The resulting t-values from all electrode pairs were modeled for group-level analyses using mixed-effects regression (see Statistical Analysis below).

#### 2.9.1 Ripple-triggered Power Analyses

Modulation of OFC activity by temporal vicinity to hippocampal ripples was evaluated in two ways: (1) Testing changes in the full OFC spectrogram (Figure 3Bi): For each frequency and time point, t-statistics of the power amplitude difference between peri-ripple and non-ripple epochs (*see* Oscillatory Activities in the OFC) were aggregated over all electrode pairs. The group means of the t-statistics were tested against permutation t-statistics, which were obtained from data that were jittered in time-dimension. For statistical thresholding, we used the 95^th^ percentile of the values and cluster sizes in the permutated data. (2) To directly test the modulation of OFC gamma-power (Figure 3Bii), we calculated a t-statistic comparing ripple-locked gamma-power to gamma-power locked to an equal number of non-ripple segments in every session.

Statistical comparisons were conducted in MATLAB, including linear mixed models (LinearMixedModel class, *fitlme*()). Our mixed models included fixed effects of ripple-occurrence, event-type, and block as well as a random subject intercept.

## 3 Results

### 3.1 One-shot learning of hidden goal location in a virtual maze

To study the dynamic modulation of OFC gamma oscillations by hippocampal activities and computational demands, we recorded local field potentials (LFP) simultaneously from hippocampal and orbitofrontal locations (Figure 1A, Figure S1A) in patients implanted with intracranial electrodes for clinical reasons (Methods, N=10 patients, see demographics in Table S1), while they performed **a goal-directed task** (Figure 1B) thought to rely on both feedforward and feedback interactions between the OFC and hippocampus (Buzsáki & Wang, 2012a; Siegel et al., 2012).

We employed the Maze Search Task (MST), in which patients navigated a series of partially observable, two-dimensional grid-world mazes (Kryven et al., 2024) (Methods). The objective of a participant in this task is to minimize the distance traveled to a hidden exit, which is located in different places in different mazes (Figure 1B). Mazes were structured with walls and paths, and importantly, goal locations were initially occluded and became revealed part by part as the virtual agent they controlled approached the unrevealed section. This task feature kept the goal hidden until the agent was close to the goal location (see example -goal reveal, Figure 1B).

Patients played 2-3 blocks, each comprising the same 30 unique mazes, which were presented once per block in a randomized order. In this manuscript, we analyzed data from the first two blocks. The mazes were set in grids of varying dimensions (e.g., 4x6, 8x7). The agent’s initial location was always the northwest corner of the maze, and the goal location varied across mazes (Kryven et al., 2024). Participants completed 67% of mazes with equal or fewer steps on block 2 (Figure 1C, Wilcoxon signed rank test - P = 0.002, also see Figure S2A for participant-specific breakdown). This provides behavioral evidence consistent with rapid learning; however, our current definition does not benchmark improvement against maze-wise null models; thus, exceedance over chance or structural affordances cannot be fully established.

We hypothesized that Reaction time (RT) would be longer in meaningful navigational events compared to when the agent simply moves along a linear open path (‘Open Path’). We also assumed RT would shorten with learning (Cotton & Ricker, 2022). We also assumed RT would shorten with learning (Cotton & Ricker, 2022). To test this hypothesis, separate linear mixed-effect model regression was estimated for each event type of interest with a random effect of Subject (Gordon et al., 2014), Methods). When we aggregated all trials, we found that successful learning is a dominant modulator of RT: Successful learning (success), block, and the interaction between them were defined as main predictors, and the subject variable was included as random intercepts (RT ∼ success + block + success*block + (1|subj)). P-values for all main predicators were < 0.003. We then focused the analysis on successful trials - Event type, block, and the interaction between event type and block were included as main predictors, and the subject variable was included as random intercepts (RT ∼ event type + block + event type*block + (1|subj)). The regression models revealed that participants were significantly slower when making the first move in a new maze (‘Maze outset’) compared to responses in Open Path or at bifurcation points (‘Choice Point’) (Figure S2B). These differences suggest that patients deliberated in maze locations that required planning, decision-making, and storing information for future trials, more than in open-path locations in which only one movement option was possible. We did not observe a change in RT across blocks in either Pre-goal locations (before the goal was revealed) or open-path locations (Figure 1D). However, without a matched second-block control of novel mazes, practice and strategy changes across blocks cannot be fully excluded as contributors to equal/improved step counts. All RT differences were statistically significant after correcting for multiple comparisons (p < .05/4). For RT of successfully learned mazes, there was no significant main effect of block or interaction effects between block and event type (Methods, all P > .1).

To pinpoint differences between 1st and 2nd traversals, we separated subsequent analysis between behavioral locations in the maze (event-types, Figure 1B). In the subsequent electrophysiological analysis, we used open-path behavioral-events as a baseline state that reflects cognitive demand common to all behavioral events in the maze, such as processing of visual input and motor planning. We used that baseline to contrast behavioral events, such as maze outset and around goal-reveal, in which memory processes are involved. We focused on two computationally critical time points in terms of figuring out the quickest way out of the maze, in which several models would hypothesize increased interactions between the executive processes and the memory system (Heckhausen & Gollwitzer, 1987; O’Reilly, 2020): First - maze outset, in which participants pull up previous experience to plan maze traversal, and pre-goal reveal, in which they are close to completing their aim, but the goal is not revealed yet. In the pre-goal locations (i.e., the salient goal is not lit up yet), the visual input is similar to ‘open path’ locations, in which movement direction is chosen based on the previously chosen path. Additionally, based on behavior markers - from the three goal-related sub-positions (Figure 1B – 2^nd^ row) - pre-goal locations had similar reaction times to open-path locations (Figure 1D), allowing us to use ‘open-path’ positions as a reference to remove movement-related activity from our neurophysiological analysis.

### 3.2 OFC Gamma power is modulated by memory demands

A core feature of orbitofrontal cortex (OFC) function is thought to be predicting specific outcomes that should follow sensory events or behavioral choices (Rudebeck & Murray, 2014). As such, OFC is thought to perform critical computations during spatial memory tasks by linking cues and spatial features with goal-relevant choices and desired outcomes (Farzanfar et al., 2023; Wikenheiser & Schoenbaum, 2016).

iEEG gamma oscillations (50-100Hz, Methods) have been associated with enhanced neural activity reflecting underlying neocortical computation (Buzsáki & Wang, 2012b). We hypothesized behavioral events associated with key cognitive computations would be associated with an increase in gamma power. Specifically, we predicted that gamma power would increase during the outset of the maze, when participants would be surveying the maze and planning, and when discovering the goal, when associations between maze-layout and goal-location could be formed for use on the subsequent trials. We also hypothesized that neural processing at these moments would differ depending on whether participants were exploring and learning the maze for the first time, as compared to when the maze was repeated and they could rely on memory to select optimal paths. Thus, our analyses compared the effect of high-computational load (maze outset and goal reach) vs low computational load (open-path) and first- and second-traversal of the maze set.

We analyzed activity in OFC channels during trials in which successful learning took place (Figure S1, Table S3, Methods). To characterize changes in the gamma band, we calculated time-frequency responses (TFR), which allow identification of spectral changes with a high temporal resolution, time-locked for each behavioral time point, relative to the preceding period (1-sec baseline, Methods). We separated this analysis between the first and 2nd trials to reveal differences due to experience (Figure 2A, Methods). To quantify changes in OFC activity, we calculated the “gamma-change percentage” (GCP) as the average of TFR values in the (50-100Hz range during a [0.15-0.35] sec time-window locked to a behavioral event.

We conducted a factorial analysis on all GCP values aggregated from high computational load behavioral time points (maze onset, pre-goal, and goal reveal, see Figure 1B) vs. low computational load (open path, at least 2 steps away from the goal). We asked whether the behavior and experience (block) modulate GCP on OFC channels during successful trials. A mixed-effects linear model (*lme*(), Matlab) was used to ask whether different behavioral positions in the maze (event types) and experience (block 1/2) modulate GCP. A random intercept for each participant was included to account for repeated measures. There was a significant main effect of event types, indicating that GCP was higher after high computational load behavioral events than open-path behavioral-events (P < 10^-20^, F(3,18691) = 36.6). A significant main effect of experience was also observed (P = 0.003, F(1,18691) = 8.29). Importantly, there was a significant block × position interaction (F(3,18691) = 11.37, P < 10^-6^) indicating that gamma activity at different behavioral-events differed between exploration and memory-guided trials.

We then asked whether this interaction manifested in different dynamics during learning at high-load behavioral time points. To that end, we separated this analysis for each high-load behavioral timepoint, comparing each one against the open path trials - see Figure S3 for a detailed bootstrapping analysis for each high-load timepoint vs open path position. For visualization purposes, we also conducted non-parametric statistical analysis beyond the factorial analysis.

We first analyzed activity related to maze-onset and found a significant increase in gamma-power during maze-onset in both blocks (Wilcoxon signed-rank test, P = 0.0009, 0.001 for blocks 1 and 2, respectively, Figure 2B-C), in line with OFC’s suggested role in planning the to-be-traversed path during goal-directed decision making (Rudebeck & Murray, 2014; Wikenheiser & Schoenbaum, 2016).

We next analyzed activity related to the goal-reveal period – the moment participants can first learn the goal location for the current map context on exploration trials and confirm its location on experience trials. OFC’s proposed role in encoding the identity and subjective value of goals, such as rewards, might predict increased computations during value update (Rudebeck & Murray, 2014; Wallis & Kennerley, 2010), which is expected in goal-reveal locations. Based on this prediction, we expected to find increased activity on goal-reveal timepoints in both blocks, but we only found a significant increase in this marker following goal-reveal in the 2nd trial (Wilcoxon signed-rank test, P = 0.5 for exploration block, Figure 2B, and 0.004 for experience-based block, Figure 2B-C). This could suggest that the OFC’s involvement increases during memory-based trials relative to exploration trials.

When a new maze is revealed, planning the optimal path towards the goal (exit) would include retrieval of previous experience in the 2^nd^ trial, but not in the 1^st^ exposure. Contrasting these two scenarios is a way to reveal activity related to retrieval functions. We expected that exploration in the first trial might be dominated by the executive functions, whereas experience-based planning would involve retrieval processes, i.e., mental search through previously learned mazes. We hypothesized that memory retrieval would introduce a lag in OFC dynamics in the second block, which was not present in the first exploration block. To test this hypothesis, we estimated the lag between block-1 and block-2 gamma power time course. To that end, we calculated a gamma-time-course, which is the averaged TFR gamma-range, time-locked to entering the same behavioral location (Figure 2A, gamma time-courses are plotted above the bottom panels). We used the amplitude cross-correlation method (Methods) calculated between block-1 and block-2 time-courses. A negative lag would indicate that block 2 gamma-power peaks earlier than in block 1. We found a significant cross-correlation in both the planning phase and the goal-reveal, but not in open path locations (Figure 2D, plots using *xcorr*(), Matlab, *corrcoef*(): R=0.24,0.27,0.01 and P = 10^-19^, 10^-25^, 0.52 for maze onset, goal-reveal, and open path, respectively). In line with our prediction, experience-based planning was associated with a *delay* in the peaking of activity in block 2 relative to the first block, whereas the opposite relationship was found at goal-reveal locations (Figure 2D).

Based on a body of work showing anticipatory OFC activity coding for expected outcomes (Howard & Kahnt, 2021; Wikenheiser & Schoenbaum, 2016), we hypothesized that OFC activity would reflect anticipation of the goal locations on experience-based trials. We analyzed GCP activity on OFC channels at “pre-goal” locations, when the virtual agent was located one step away from the goal reveal (Figure 2E). To characterize how proximity to goal location modulates gamma power, we first compared open-path locations in the maze (constrained by maze structure that were further than 2 steps from the goal) to pre-goal locations (one step away from goal reveal). Note that different spatial positions in different mazes are associated with the same behavioral event, i.e., open-path or pre-goal/goal location. We will refer to these behavioral events as event types in the maze, and they occur at variable spatial locations on the screen. We repeated the factorial analysis, including pre-goal locations and open path locations, and asked whether event type and experience modulate gamma-power. There was a significant main effect of experience (Block 1/2, P = 0.004, F(1,13962) = 8.03) that was qualified by a significant block × event type (pre-goal vs open path) interaction (F(1,13962) = 9.4, P 0.002). Follow-up analyses revealed that, on experience-based trials (block 2), gamma-power decreased in the pre-goal locations relative to the open-path locations (Figure 2F). In contrast, there were no significant differences between the two trial types in the first (exploration) block. Furthermore, when contrasting the pre-goal locations in the 2^nd^ block with the 1^st^ block trials, we discovered a decrease in gamma-power in these behavioral events with experience (Figure 2G, Wilcoxon signed-rank test, P = 0.006).

Overall, our results indicate that OFC gamma power is modulated during events that involve prospectively planning routes, anticipating goals, and confirming goal locations, and by experience (i.e., the difference between exploration and memory-guided performance).

**Figure 2.**
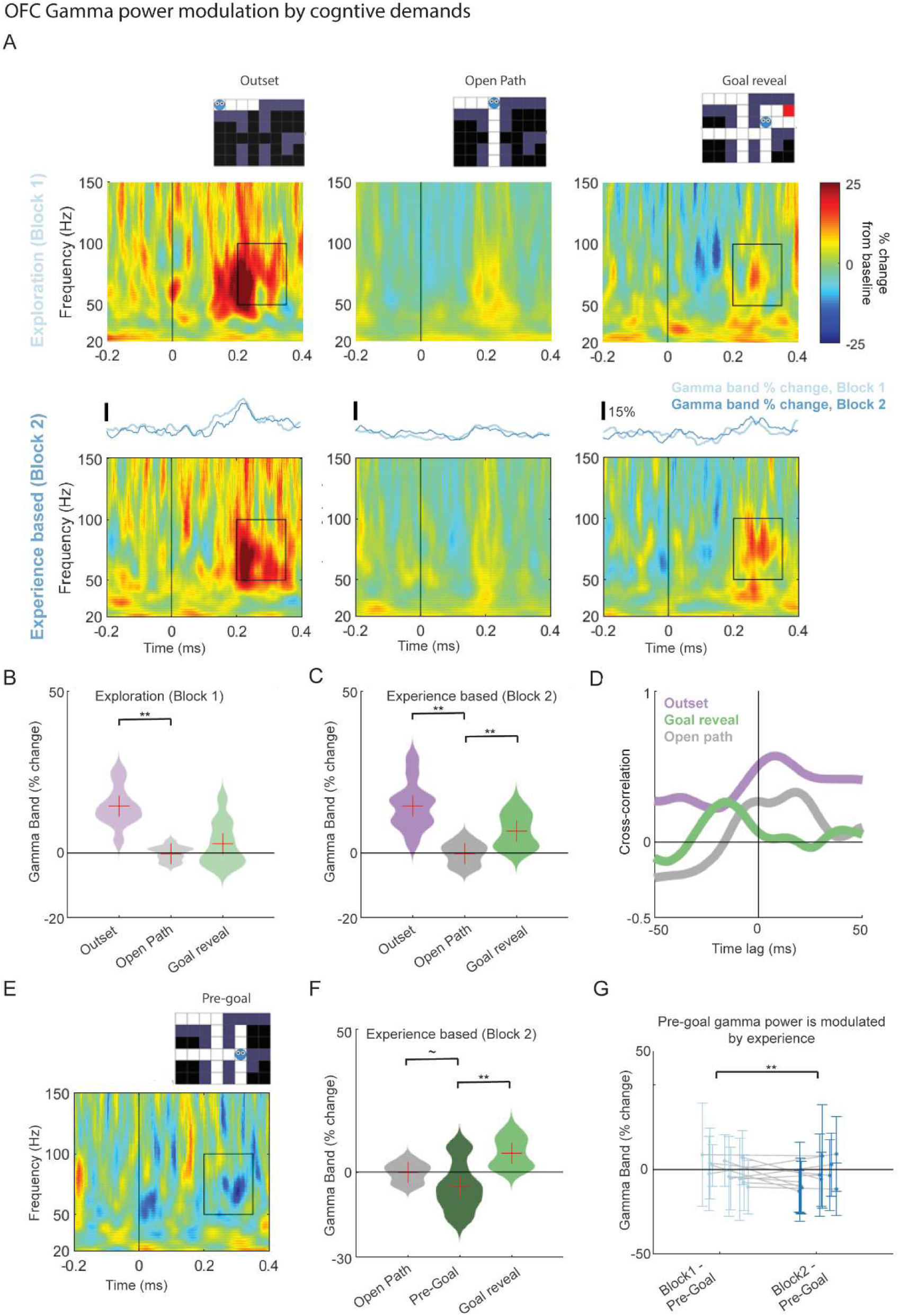
OFC gamma-power is modulated by memory demands during successful learning. (**A**) Averaged time-frequency responses (TFR) were used to identify rapid changes in gamma-power. Top row – TFR of first block trials (exploration); Bottom row - second block trials (experience-based). OFC channels demonstrate a significant increase in the gamma-band (50-100 Hz) following maze outset (left column - top and bottom) and during goal-reveal in the second block (right column, bottom), but not during the first exploration (right column, top). TFRs for open-path locations (middle column) do not exhibit significant power modulation. Color bar indicates % change from the preceding 0.5 sec periods, with voxels over max/min values saturated. The waveforms between TFRs illustrate average gamma-band values for each time point: a lighter time course for the exploration block and a darker time course for the experience-based block. (**B, C**) Population data showing gamma power modulation by behavioral location with varying cognitive demands during exploration (B) and memory-based trials (C). Gamma power increased after maze onset in both blocks (purple distribution), with a significant increase upon goal reach in the second block (green distribution) but not in the first. All comparisons are relative to open-path locations (gray distribution). Each violin plot represents the distribution of sums of TFR windows marked in panel A ([0.15 0.35]sec, 50-100Hz) for OFC channels (N=11 channels). (**D**) Cross-correlation of gamma-power time-courses of block 1 and block 2 (average time course plotted on panel A) demonstrates a delayed time course during the outset and open-path TFRs but not for the goal location. (**E**) While goal-reveal locations exhibited an increase of gamma-power in the second block (A), gamma-power **decreased** in pre-goal locations (one step before goal reveal). TFR definitions are described in panel A. (**F**) Population comparison of pre-goal location gamma power (time/frequency window as in B) in the second block, to open-path locations (P = 0.057) and goal-reveal locations (P = 0.0003), (N=11 channels). (**G**) Population comparison of pre-goal location gamma power (time/frequency window as in B) in the second block, relative to the exploration phase (ii, N=11 channels). Sign rank test, P = 0.01.

### 3.3 Ripple-locked orbitofrontal gamma-power is associated with subsequent memory performance

Goal-directed navigation is likely to engage coordinated activity across the hippocampus and OFC (Wikenheiser & Schoenbaum, 2016). One mechanism that could account for coordinated activity is increased OFC activity time-locked to hippocampal ripples during navigational events. The hypothesis we went out to explore is whether time-points that were associated with increases in OFC gamma-activity would also be associated with heightened synchronization between cortical patches during hippocampal ripples (Buzsáki & Wang, 2012b). To test this hypothesis, we analyzed data from a subset of patients in which intracranial recordings from both the medial temporal lobe (MTL) and OFC were available (Methods, N=11 electrode pairs from 7 patients/sessions, Figure 1A, left). The majority of MTL contacts were placed in the hippocampus, but we also assessed adjacent contacts in the amygdala and parahippocampal gyrus (Figure S1, Table S3), given that ripples have been observed in both areas (Clemens et al., 2007; Geva-Sagiv et al., 2023; Helfrich et al., 2019). We collectively refer to this group as ‘MTL’ electrodes. To examine the presence of fast-frequency ripples in MTL iEEG recordings, we used a recently developed algorithm to isolate these events in the human hippocampus and parahippocampal cortex (Figure S1, Methods (Liu et al., 2022; Norman et al., 2019, p. 201, 2023)). We did not find statistically significant modulation of ripple rate around behavioral events in the maze (Methods, Figure S4A).

First, to check whether OFC oscillatory activity is modulated by the presence of MTL ripples, we compared the spectral profile of OFC channels during ripples to their profile during non-ripple segments (Methods). The analysis revealed significant differences in spectral profile time-locked to MTL ripples (Methods, Figure 3Bi). We found that OFC-gamma power significantly increased during ripple times relative to non-ripple windows (Figure 3Bii, Wilcoxon signed-rank test P = 0.027). To test whether this difference is specific to the OFC, we repeated these analyses in gray-matter (GM) contacts residing outside the OFC (Figure S4B, Methods) and did not find any changes in spectral profile during ripple times (Figure S4C, inset) nor did we find a significant change in gamma-power (P=0.31) during ripples in GM electrodes in non-OFC regions, suggesting anatomical specificity for OFC region.

Next, we asked whether ripple-locked OFC gamma power is modulated during successful learning. To that end, we focused on successfully learned trials and found that the ripple-locked OFC gamma power peaks at maze outset in both blocks and goal reveal at the exploration phase, relative to lower-computational load events (open path and goal entry, Figure 3A). To statistically test whether gamma power was modulated by changes in computational load during learning of the task, and whether this modulation is specific to ripple times, we fitted a linear mixed-effect model with a random effect of Subject (Gordon et al., 2014), testing for effects of event-type, experience (block), and ripple-vicinity on OFC gamma-power as well as the interaction between ripple-vicinity and the behavior-related main effects. A mixed effect model demonstrated a significant effect of ripple-vicinity and event-type and a significant interaction between block-type and ripple vicinity (ripple-vicinity – F(1) = 8.5, P = 0.003; event type - F (8) = 5.1, P < 10^-5^; block-type = F(1) = 5.2, P = 0.002, block*ripple-vicinity – F(1) = 5.8, P = 0.01). To unpack the effect of experience and understand whether it is driven by specific event types, we calculated the effect size for each behavioral event separately (Figure S5A). We found that ripple-locked OFC gamma power decreased in the second block at time points that are associated with plan execution (open path and pre-goal reveal), but that the ripple-locked OFC gamma power during high computational load events (maze-outset and goal reveal) was not significantly modulated by experience (Figure 3C inset, Figure S5A). This suggests that gamma power is strongly modulated by ripple-vicinity and event-type, and that learning effects are dependent on the cognitive demands of behavioral events.

Finally, we asked whether ripple-locked OFC gamma power is related to successful learning. In the current learning paradigm, we hypothesized that ripple-locked OFC gamma activity on exploration trials (1^st^ block) would be indicative of performance on the 2^nd^ attempt (block 2). To test this hypothesis, we added to the analysis block-1 trials from unsuccessful (degraded) trials. We found that, during block 1, ripple-locked OFC gamma power at maze-onset was higher for successfully learned mazes than for degraded performance mazes (Figure 3D, inset). To statistically test whether ripple-locked OFC-gamma power on block 1 is related to learning, we fitted an additional mixed-effect model with a random effect of Subject, testing for effects of event-type and OFC ripple-locked gamma power during the exploration block, on subsequent performance in the second block. This model revealed a main effect of the event-type and a significant interaction between gamma power and event-type in the first block trial on subsequent performance in block 2 (event – F(8) = 11.1, P < 10^-12^, interaction between gamma-power and behavioral-event - F = 3.06(8), P = 0.02). To unpack this effect for different event-types, we compared ripple-locked OFC gamma power during the first block for different event-types, for successfully learned mazes against degraded ones. We found that, at block-1 maze-onset and pre-goal locations, ripple-locked OFC gamma power was higher for successfully learned mazes than for degraded performance mazes (Figure 3D, inset, Wilcoxon signed-rank test, P = 0.001 for Maze-onset locations and P = 0.02 preceding goal reveal, see also Figure S5B for effect size). These results demonstrate that ripple-locked OFC gamma power is selectively enhanced during high computational load events and predicts successful learning, with stronger coupling during initial exploration distinguishing successfully learned from degraded performance trials.

**Figure 3.**
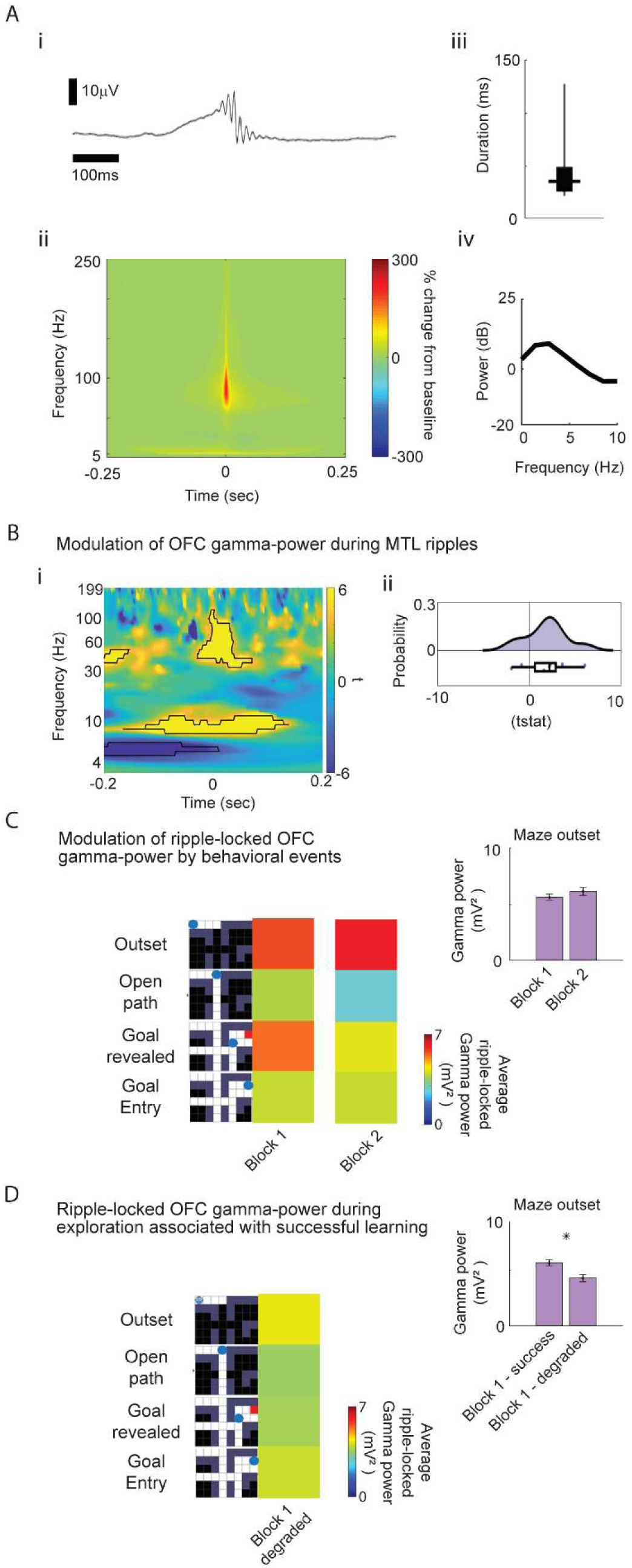
OFC gamma power is modulated by both high-frequency ripple activity and behavior. (**A**) Ripples were detected from MTL LFP recordings (anatomical locations on Figure S1 and Table S1): (**i**) Grand average of unfiltered iEEG aligned to the maximum ripple peak (mean ± s.e.m., n = 12812 ripple detections). (**ii**) The average ripple-peak-locked time-frequency response (percentage change from 1-s baseline) highlights the band-limited frequency profile of detected ripples. (iii) Box plot of ripple duration (mean ± SEM: 42.4 ± 0.2 ms). (**iv**) A spectral analysis of the grand average iEEG (calculated from −1 s to +1 s using FieldTrip’s *mtmfft*() function for frequencies from 1–10 Hz in steps of 0.5 Hz) revealed a spectral peak at 3 Hz. **(B)** Modulation of OFC oscillatory signatures during hippocampal-area ripples: (i) OFC time-frequency analysis around hippocampal ripples (Methods) - Color represents the t-stat of the oscillatory power compared against the values from data generated by time-frequency analysis of non-ripple segments. (ii) Ripple-locked OFC gamma-power (Methods) is significantly higher than non-ripple time windows. The X-axis depicts t-statistics comparing ripple-locked PAC to non-ripple segments. X-axis depicts the probability of tstat values across cross-brain couples. Wilcoxon signed-rank test : P = 0.027. (**C**) Ripple-locked OFC gamma power is modulated by both event-type in the maze and experience. Qualitative description: In block one trials, ripple-locked OFC gamma power is highest during Maze-onset (planning path to goal) and goal-reveal. Main panel - Every colorful square depicts average ripple-locked gamma-power around ripples occurring in the vicinity of event-types illustrated on the left – on block 1 (middle column) and block 2 (right column). Sub-panel – no significant difference between average ripple-locked gamma power on maze onset between blocks. (**D**) Ripple-locked OFC gamma power on block 1 is associated with successful learning. Main panel - Colors as in panel C depict average ripple-locked OFC gamma-power during exploration trials of degraded mazes (Methods). Sub-panel – average ripple-locked gamma power on maze onset on exploration trials resulting in degraded performance (degraded) is significantly lower than that of successfully learned paths (impv, left bar is same value shown in C-sub panel). Error bars depict SEM, and ‘*’ indicates significance on a Wilcoxon rank-sum test, which was added for visualization purposes. Fitted linear mixed effects models are described in text.

## 4 Discussion

In this paper, we study the role of hippocampo-orbitofrontal interactions in continuous behavior unfolding across extended time-scales, which leverages exploration, learning, and memory. To do this, we’ve used intracranial recordings from OFC and the hippocampus in epilepsy patients as they performed a 2D goal-seeking spatial task. The task proceeded in two consecutive blocks. In the first block, the task demands were primarily driven by planning and exploration, and in the second block, required memory-guided decision-making. We measured hippocampo-orbitofrontal interactions in terms of OFC gamma power functionally coupled to hippocampal ripples, finding that task-driven interactions between these areas followed a phasic, rather than a continuous nature. The hippocampo-orbitofrontal interactions were particularly prominent at the time of maze-onset and when reaching the goal during the memory-guided phase - suggesting that OFC gamma power was modulated by both learning and cognitive demands. We also found that ripple-triggered gamma increases were associated with improved behavioral performance during the memory-guided phase of the navigation, suggesting OFC-hippocampal interactions support one-shot learning and memory-guided action.

The maze paradigm introduced here requires subjects to learn a context-specific goal that must be achieved after multiple decisions, akin to navigating a decision tree in order to obtain a desired outcome. The goal of this task was not to investigate spatial navigation in the literal sense, as is done in recordings of neural activity during navigation of virtual reality mazes or mazes in physical space (Reggente et al., 2018). In this task, we investigated navigation in the more abstract sense, in which exploration and memory-based planning are used to guide a series of actions to reach context-specific reward locations. On the first encounter with each maze, patients needed to discover the goal location through exploration, but on repetition, they could retrieve past experiences to more efficiently navigate to the goal. The design of the current task allowed us to identify, with high temporal precision, timepoints in which subjects generate or retrieve plans (maze-outset) and encode or predict the locations of goals (goal-reveal) and examine distinct neural signatures of each event. However, as in other navigation-emulating paradigms, we do not have full power to dissociate between planning and memory-guided decisions. For example, people who have not learned the goal location and those who have may have made different assumptions about goal placement or task rules.

Our first finding demonstrates a modulation of OFC gamma band activity by behavioral demands of the position in the maze (event type), by experience (blocks 1-2). OFC gamma power increased during the initiation of maze navigation and goal attainment relative to open-path locations in the virtual maze (Figure 2A). These findings support the idea that, in addition to its role in memory retrieval, the prefrontal cortex may support the integration of new memories into pre-existing knowledge (Preston et al., 2004). Also, these results are in line with the idea that OFC computations peak at critical events that involve planning and linking choices and outcomes (O’Reilly, 2020; O’Reilly et al., 2014). Separating different event-types in a continuous task was key to identifying the different patterns of neural activity related to different cognitive demands of these behavioral events. We note that Maze onset and goal reveal co-occur with movement initiation and visual orienting, which may not be fully accounted for by using open-path events as controls, and our experimental design did not allow modeling motor/oculomotor covariates in the reported high-frequency analyses. The specificity of gamma-power increase in event types associated with the higher cognitive load of memory-related computations is in line with theoretical models associating prefrontal gamma-band modulation with synchronization of neural networks needed for efficient memory encoding and retrieval (Buzsáki & Draguhn, 2004; Buzsáki & Wang, 2012b).

Our second key finding was a coupling between OFC gamma activity and high-frequency ripples (Figure 3B). This coupling was cognitively selective—preferentially enhanced during high-demand events—and behaviorally predictive, with gamma power during initial maze exploration forecasting subsequent learning success. Our conservative ripple detection procedures ruled out contamination by pathological epileptic activity (Liu et al., 2022, Methods), and concurrent recordings highlighted the anatomical and temporal specificity of this synchronization.

Notably, functional coupling between hippocampus and OFC was the strongest during the planning epochs, when participants computed multi-step routes. This activity was modulated by the computational demands of each planning step rather than being uniform throughout navigation. this finding aligns with the event-based theories of hippocampo-cortical interactions, and are supported by behavioral studies that demonstrate distinct behavioral phases through analysis of eye movements (Zhao & Marquez, 2013) and reaction times (Kryven et al., 2024). Our findings agree with behavioral and computational perspectives on planning and exploration, which model cognitive computation as associated with discrete events (Balaguer et al., 2016; Kryven et al., 2024) (Balaguer et al., 2016; Callaway et al., 2022; Hirtle & Jonides, 1985; Kosslyn et al., 1974; Kryven et al., 2024). Discrete planning computations are evident in a tendency to plan paths between regions, before planning sub-paths within each region (Balaguer et al., 2016; Wang & Brockmole, 2003; Wiener & Mallot, 2003), and in increased reaction times when switching between hierarchy levels during plan execution (Balaguer et al., 2016; Kryven et al., 2024). Behaviour-theoretical models treat the events of encountering a novel environment and discovering goal-related information as particularly critical to forming and updating cognitive maps (Ho et al., 2022; Sharma et al., 2022). They further dissociate between planning computations occurring when a novel environment is encountered (Kryven et al, 2024, Zhao & Marquez 2013, Sharma et al 2022), and memory-based computations driven by previous experience (Gershman & Daw, 2017, Kool et al 2017 PMID 28731839). The correlation between ripple-locked OFC gamma changes and subsequent spatial memory performance emphasizes the pivotal role of hippocampal ripples in orchestrating information exchange for rapid learning and planning. This finding aligns with animal studies suggesting these interactions reflect coordination during hippocampal replay associated with planning (Jadhav et al., 2012; Ólafsdóttir et al., 2015, 2018; Pfeiffer & Foster, 2013) and supports the notion that hippocampal ripples provide a critical window for hippocampal-cortical communication, facilitating goal-directed behavior.

Ripples were detected in the hippocampus, amygdala, and parahippocampal cortex and pooled as MTL events for analyses to maximize statistical power, as deep brain recordings combined with cognitive behavior are rare and electrode-location is determined by clinical needs alone. Because ripple events were pooled across hippocampus, parahippocampal gyrus, and amygdala, these analyses cannot isolate hippocampal-only contributions to OFC coupling. This may also explain why our data does not reveal an increase in MTL ripple-rate around timepoints associated with encoding and retrieval (Figure S4A) as previous studies have shown for strictly hippocampal ripples (Norman et al., 2019). However, other studies have found hippocampal ripple-rate to be stable across a series of associative memory tasks (Chen et al., 2021). Our study may offer a way to resolve apparent contradictions in previous reports, by suggesting that different types of memory-employing tasks would be reflected by increase of MTL-OFC coupling rather than a local increase in ripple rate.

Our findings bridge a knowledge gap that exists between animal and human research. It has been well demonstrated that MTL-ripples during post-learning rest or sleep are associated with increased hippocampal-neocortical communication that promotes memory consolidation in both rodents (Khodagholy et al., 2017; Maingret et al., 2016; Siapas & Wilson, 1998; Skelin et al., 2021) and humans (Geva-Sagiv et al., 2023; Helfrich et al., 2019; Jiang et al., 2019; Ngo et al., 2020; Sanda et al., 2021; Skelin et al., 2021). Evidence from rodent studies suggests that hippocampal ripples during wake behaviors drive cortico-hippocampal interactions that underlie goal-directed learning (Denovellis et al., 2021; Dupret et al., 2010; Fernandez-Ruiz et al., 2019; Jadhav et al., 2012; Michon et al., 2019, p. 20; Papale et al., 2016). Our findings, in which OFC gamma power synchronized to ripple events was enhanced specifically during computationally intensive phases and distinguished successful from failed learning trials, support the framework suggesting that ripples in the human hippocampus and surrounding structures offer a critical window for information exchange during critical timepoints of memory retrieval and update (which peak at planning and goal-reach in the MST game).

Our study highlights the intricate dynamics of neural synchronization in OFC and how the hippocampus and surrounding areas may coordinate this synchronized activity during high_-_frequency ripples (Overton et al., 2025). Our findings bridge gaps between human and animal research and pave the way for future investigations into the roles of high-frequency ripples in coordinating cortical activity during behavior.

## Acknowledgments

We thank the patients for their participation in this study and Ranganath lab members for discussions and comments on the early versions of the manuscript. We thank Kamin Kim for leading data curation and software development in early phases of the project and Radhika Dhanak for help in software development. This article was funded by the Office of Naval Research (Grant N00014-20-1-2578).

## Author Contributions Statement

M.G-S. played a lead role in data curation, software, formal analysis, investigation, visualization, writing-original draft. I.S conceived the research, funded the acquisition, collected all data, and had an equal role in supervision. J.J.L. funded and supervised acquisition and project administration. S.J. contributed resources, visualization, and writing-original draft; M.K. and J.T. developed the behavioral paradigm. C.R., R.O., and E.D.B. contributed to conceptualization and formal analysis. C.R. secured funding and had an equal role in supervision. All authors provided ongoing critical review of results and commented on the manuscript.

## SUPPLEMENTARY DATA

**Figure S1:**
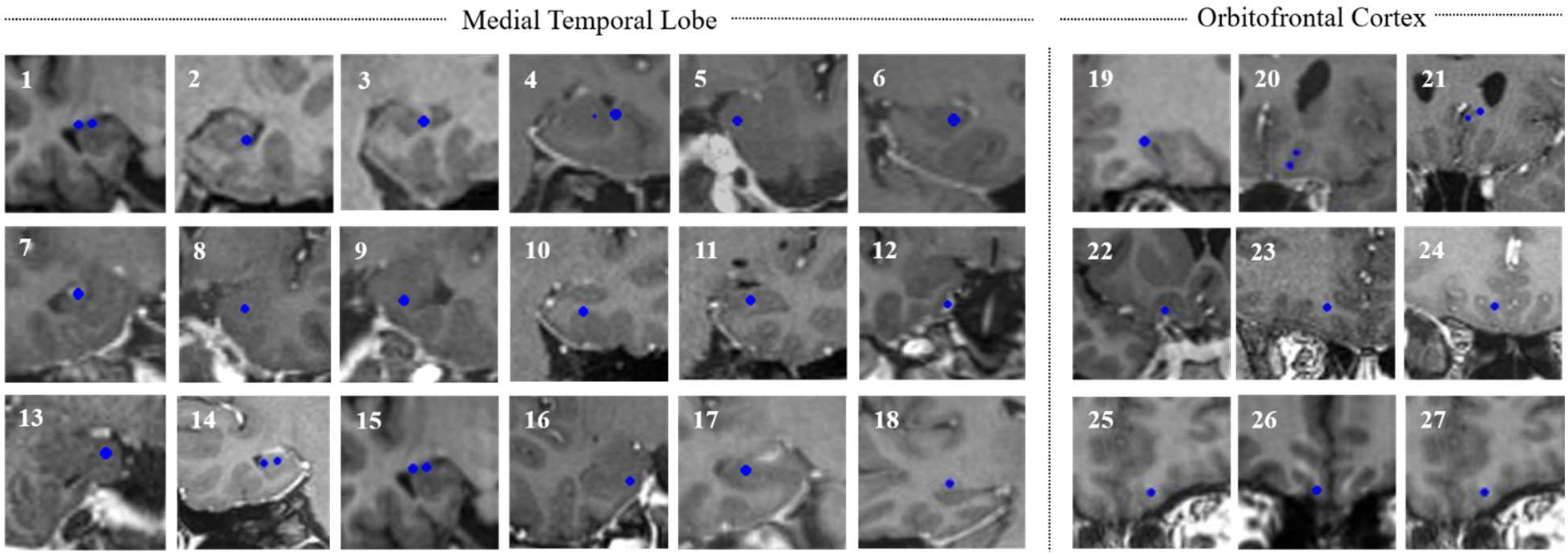
Coronal MR images for all MTL and OFC iEEG electrodes included in analysis. Blue circles depict the location of each iEEG contact on the depth electrode. See Table S3 for additional information about participants and the anatomical-location of each contact.

**Figure S2:**
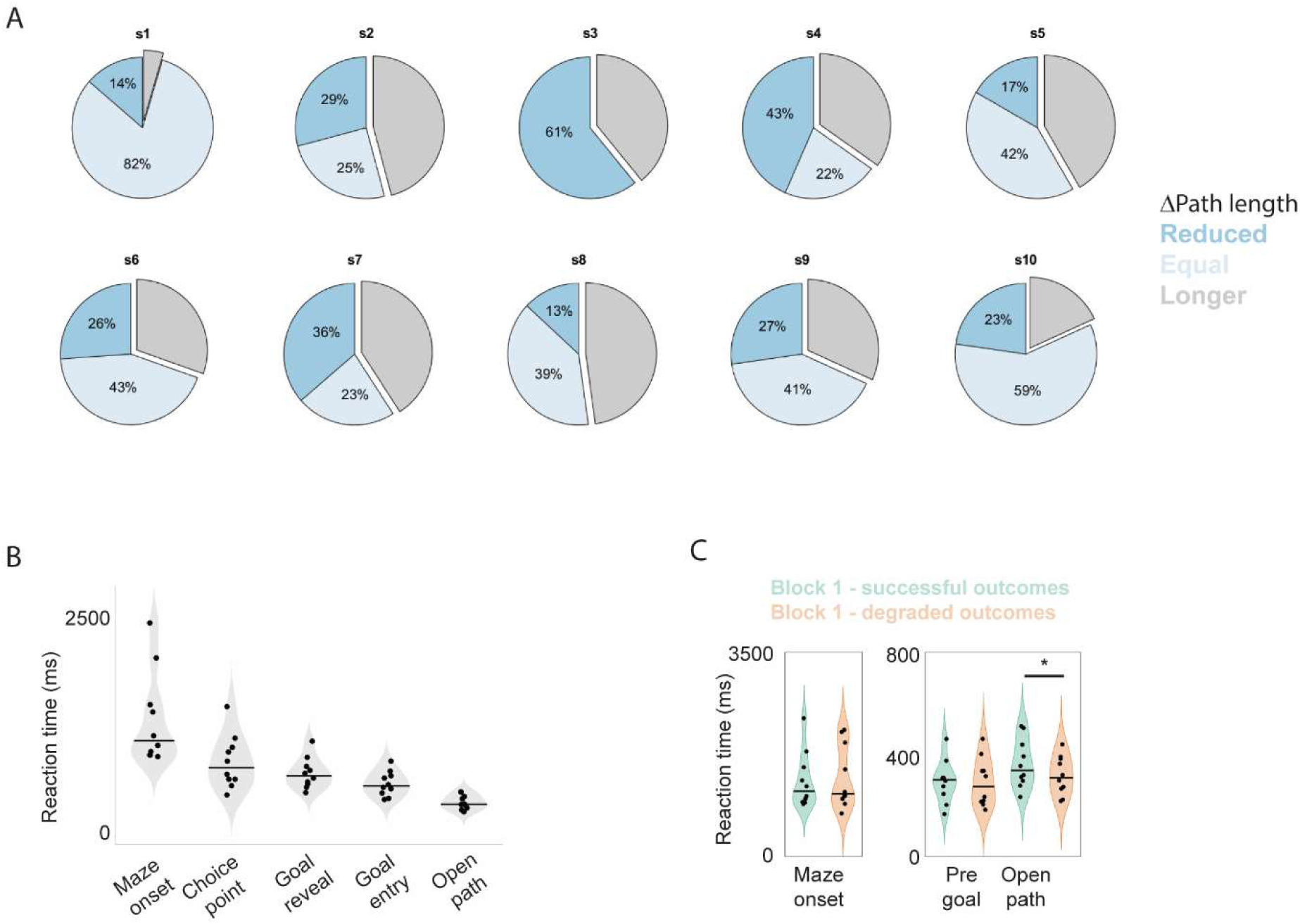
(**A**) Participant-specific performance changes over two blocks – for each participant (s1-10), each pie plot depicts the ratio of mazes that participants were able to complete with a shorter amount of steps (dark blue), with the same amount of steps (light blue), or degraded number of steps (gray). **(B)** Response time (RT) was longer at complex navigational events compared to neutral moves (i.e., Open Path). Data points depict average RT from individual participants. Horizontal lines mark group means for each event type. Compared to moving from one square to the next in an open path (Open Path), people took longer to move the agent when starting to navigate a new maze (Maze onset), when choosing one direction over others at bifurcation points (Choice Point), when the goal was revealed (Goal Reveal), and when entering a goal location (Goal Entry). (**C**) A mixed-effect linear model demonstrated that RT in the first block was modulated by the trial type (outcome successful learning vs outcome is a degraded performance; see Methods). This finding was primarily driven by differences in open-path RTs, while maze-outset and pre-goal RTs did not differ significantly between successful and degraded outcomes during the first learning block.

**Figure S3:**
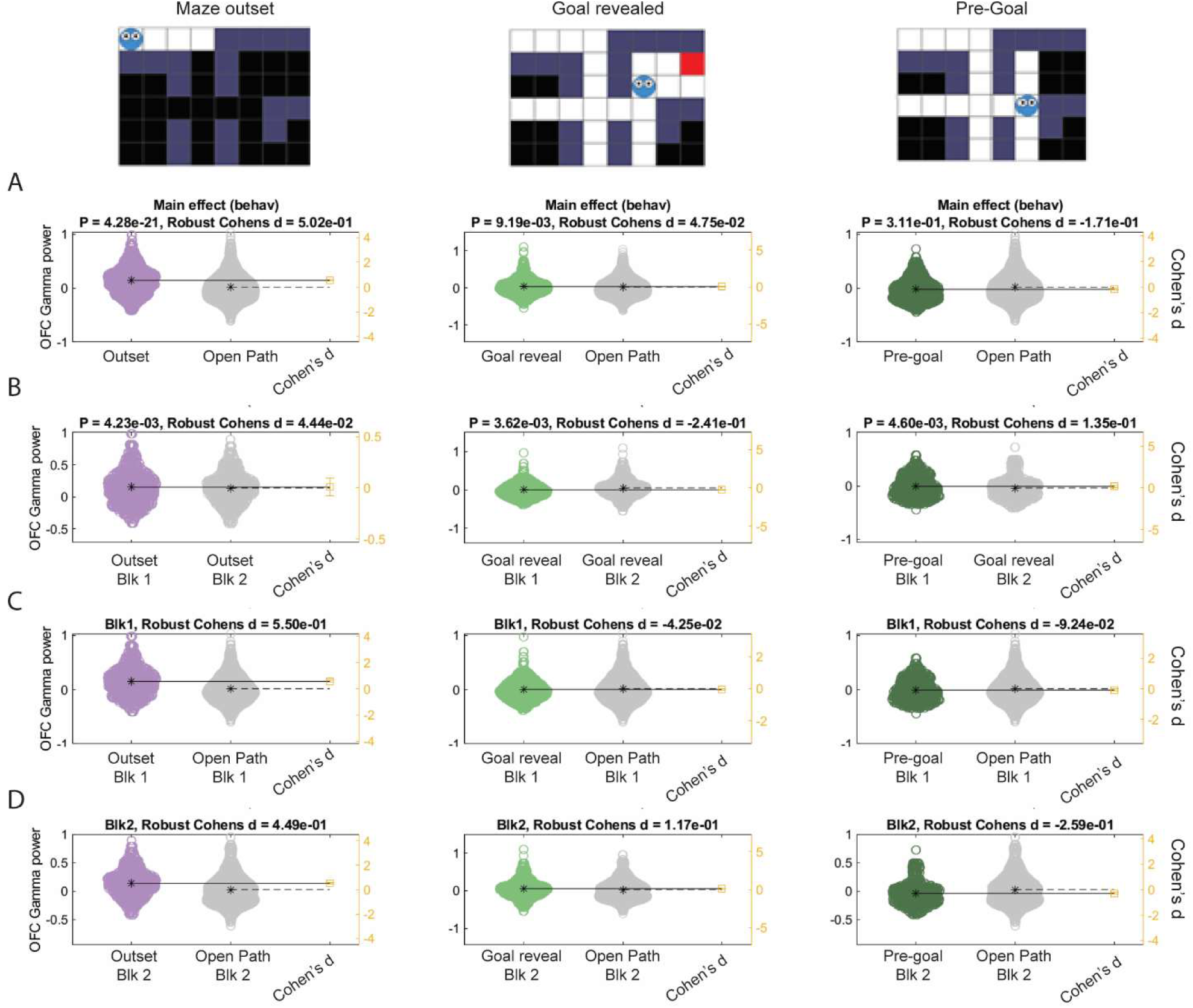
OFC Gamma power is modulated by experience and behavioral demands. We calculated the unbiased estimate of Cohen’s d (Cousineau, 2005) (Hedge’s g) for gamma-change percentage (GCP, see methods) in different behavioral locations - maze outset (leftmost column), goal reveals (middle column), and pre-goal (right column). Gardener-Aleman plots (M. J. Gardner & Altman, 1986) were generated to visualize confidence intervals. **(A)** Robust Cohen’s d is plotted for all trials from behavioral timepoint tested in each column, against the open-maze trials (low computational load). P values and Cohen’s d are plotted for each panel. **(B)** Robust Cohen’s d is plotted for all trials from the behavioral timepoint tested in each column, from block 1 (exploration) against block 2 (experience-based). P values and Cohen’s d are plotted for each panel. **(C)** Robust Cohen’s d is plotted for all trials from behavioral timepoint tested in each column, against the open-maze trials (low computational load) for block 1 only. Cohen’s d was plotted for each panel. **(D)** Robust Cohen’s d is plotted for all trials from behavioral timepoint tested in each column, against the open-maze trials (low computational load) for block 2 only. Cohen’s d was plotted for each panel. Blk X Location Interaction Effect P = 0.026, 0.0002, 0.002 for outset, goal reveal, and pre-goal, respectively.

**Figure S4:**
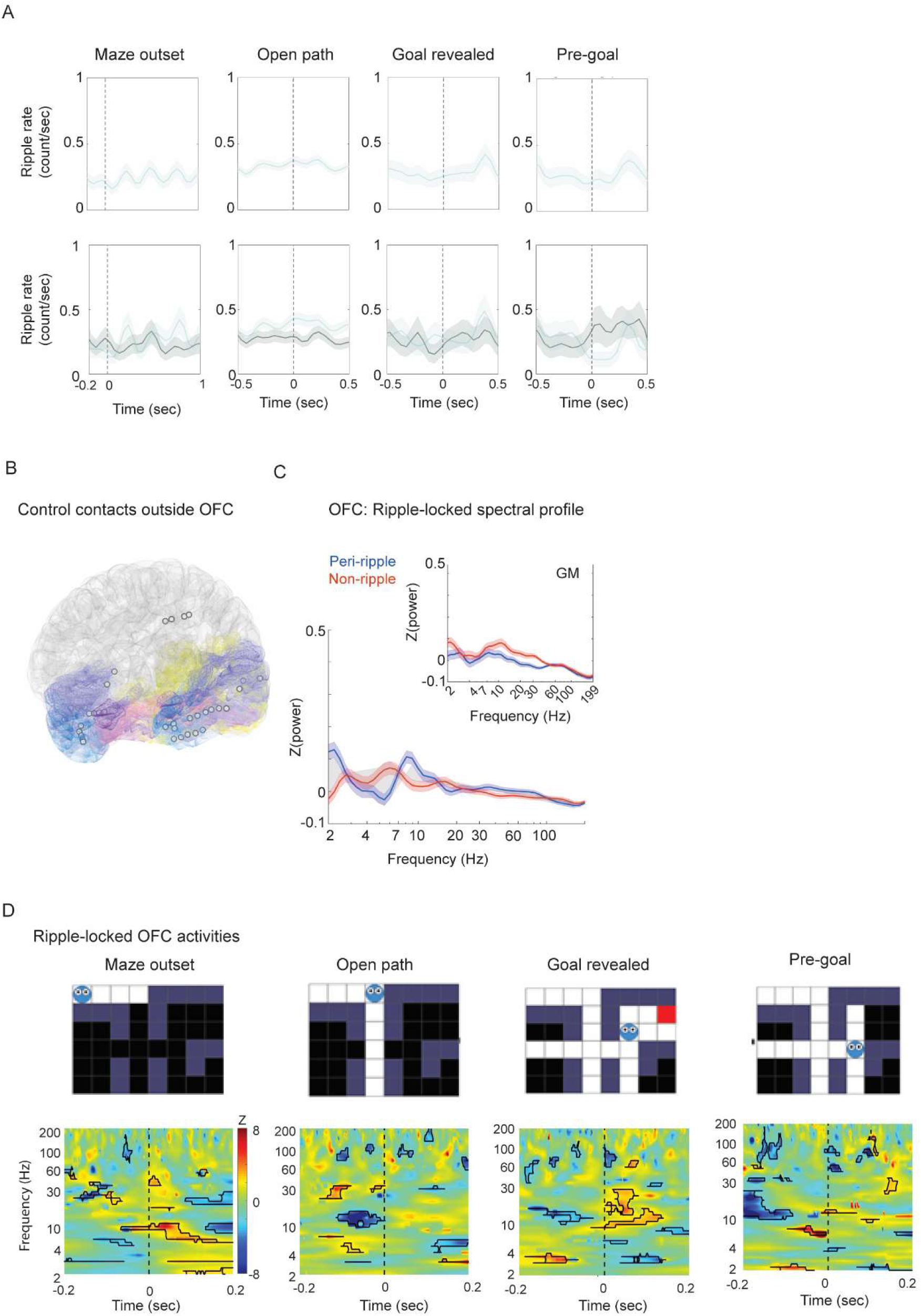
OFC oscillatory activity modulated in the vicinity of hippocampal ripples. **(A)** Ripple rate is stable around behavioral events. Traces depict the grand average of PSTH from each navigational event type. The dotted vertical lines mark the onset of the navigational event that the PSTHs are time-locked to. Ripple rates were not significantly different from the surrogate data (Methods). **(B)** iEEG contact locations are overlaid on a standard (Montreal Neurological Institute (MNI) brain template (see Methods for electrode localization) for gray-matter contacts located outside OFC that are used for control-analysis in panel C. **(C)** Average OFC spectral profile around ripples (blue, averaged over [-0.2 0.2] sec around ripple peak occurring during block 1 trials that led to improved block 2 trials), demonstrates significant differences in lower frequencies (0-11Hz) relative to non-ripple segments (red). Inset: The same analysis performed on gray-matter contacts located in other locations (see panel B) does not yield these differences and demonstrates the anatomical specificity of these changes (Methods). **(D)** OFC time-frequency analysis around hippocampal ripples for specific behavioral locations in successful 2nd block trials - Color represents the z-score of the oscillatory power compared against the values from surrogate data generated by shuffling data in the time dimension. Black outlines mark time-frequency clusters where the oscillatory activity was significantly different from surrogate data. Vertical dotted lines mark the time stamp for the peak of MTL ripples (Methods).

**Figure S5:**
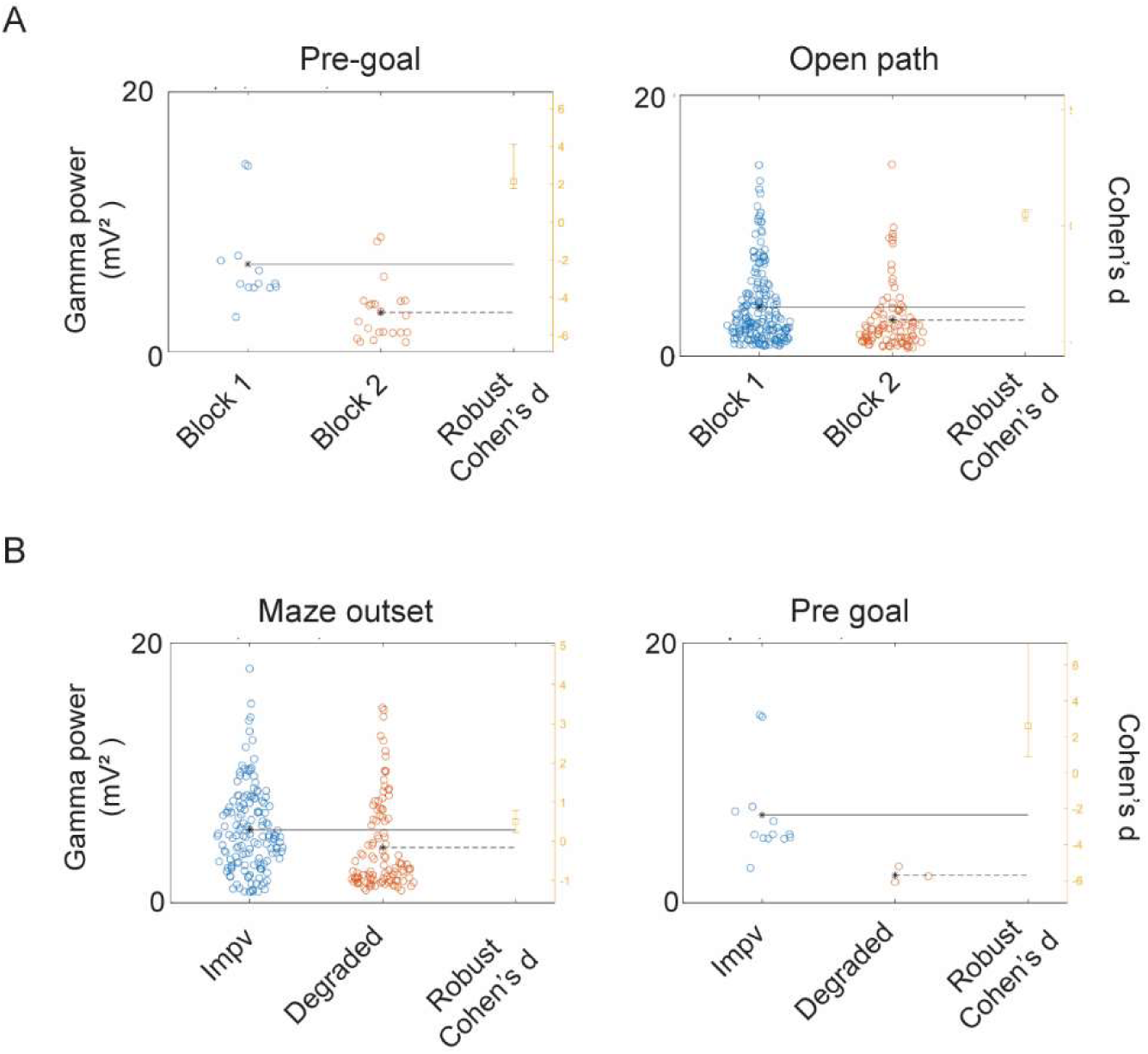
Ripple-locked OFC Gamma power - effect size calculation. **(A-b)** We calculated the unbiased estimate of Cohen’s d (Cousineau, 2005) (Hedge’s g) for ripple-locked gamma power at different behavioral locations to complement Figure 3C-D. Gardener-Aleman plots (M. J. Gardner & Altman, 1986) were generated to visualize confidence intervals. **(A)** OFC ripple-locked gamma power is reduced with learning - In successfully learned mazes - Robust Cohen’s d is plotted for all block 1 trials (blue) from behavioral timepoint tested (maze outset, left panel, and pre-goal in right panel), against block 2 trials (red). The effect size, as measured by Cohen’s d, was d = 1.29 and 0.3 for pre-goal and open-path locations, respectively, indicating a medium-large effect. **(B)** Robust Cohen’s d is plotted for all block1 trials leading to successful trials (blue markers), from behavioral timepoint tested (maze outset, left panel, and pre-goal in right panel), against trials leading to degraded performance (red markers). The effect size, as measured by Cohen’s d, was d = 0.4 and 1.3 for maze outset and pre-goal locations, respectively, indicating a medium effect.

**Table S1.** Patient demographic information, electrode count, and seizure onset zone for each patient.

| Subject | Gender | Age | Implanted Hemisphere | Seizure Onset Zone (Soz) | Handedness | MTL sites | OFC sites |
| --- | --- | --- | --- | --- | --- | --- | --- |
| Sub1 | M | 29 | L,R | L. Amygdala,<br>R. HPC, L.PHG, R. MTG | R | 3 | 1 |
| Sub2 | M | 22 | R | R. HPC | L | 3 | 0 |
| Sub3 | M | 33 | L,R | R. PHG,<br>R. Temporal white matter | R | 3 | 2 |
| Sub4 | F | 51 | R | R. Amygdala,<br>Temporal pole | L | 2 | 2 |
| Sub5 | M | 38 | L | L. PHG, L. STG | R | 1 | 1 |
| Sub6 | M | 43 | L,R | L. Cerebral white matter, R. MTG,<br>R. STG | R | 1 | 1 |
| Sub7 | F | 18 | L | L. Temporal pole,<br>L. Amygdala | R | 1 | 1 |
| Sub8 | M | 23 | L,R | - | R | 1 | 1 |
| Sub9 | M | 23 | L,R | - | R | 1 | 1 |
| Sub10 | M | 33 | L,R | R. HPC,<br>R. Amygdala | R | 2 | 1 |
| TOTAL |  | 31.3<br>(9.91) |  |  |  | 18 | 11 |

Abbreviations: R. = Right; L. = Left; HPC = hippocampus; PHG = parahippocampal gyrus; OFC = orbitofrontal cortex; STG = superior temporal gyrus; MTG = middle temporal gyrus; ITG = inferior temporal gyrus

**Table S2.** Behavioral events in the virtual maze – event names correspond to the names used in Figure 1A. See detailed description of the task in the Methods section. A trial consists of N movements across N locations. A trial starts on T_1_ and ends on T_N_ when entering the goal location.

| Behavioral in the virtual maze | Spatial position on screen | Time in trial: $T_1, \dots T_N$ | Occurrences per maze |
| --- | --- | --- | --- |
| Maze onset | Top left corner | $T_1$ | 1 |
| Open path | Variable | $T_{2:N-2}$ | Multiple |
| Decision point | Variable | $T_{2:N-2}$ | Multiple |
| Goal reveal | Variable | $T_{N-3:N-1}$ | 1 |
| Goal entry | Variable | $T_N$ | 1 |

**Table S3:** List of iEEG contacts included in analysis, IDs correspond to MR images in Figure S1. Subject corresponds to patient (sub1-10, Table S1). Locations based on Yale Brain Atlas (McGrath et al., 2022) (Methods, High-resolution cortical parcellation based on conserved brain landmarks for localization of multimodal data to the nearest centimeter). Abbreviations are as in Table S1.

| ID | Subject | Implanted Hemisphere | Location (YBA label) |
| --- | --- | --- | --- |
| 1 | Sub1 | L | HPC |
| 2 | Sub1 | R | HPC |
| 3 | Sub1 | R | HPC |
| 4 | Sub2 | R | Amygdala |
| 5 | Sub2 | R | HPC |
| 6 | Sub2 | R | HPC |
| 7 | Sub3 | L | HPC |
| 8 | Sub3 | R | HPC |
| 9 | Sub3 | R | HPC |
| 10 | Sub4 | R | HPC |
| 11 | Sub4 | R | HPC |
| 12 | Sub5 | L | PHG |
| 13 | Sub6 | L | HPC |
| 14 | Sub7 | L | HPC |
| 15 | Sub8 | L | HPC |
| 16 | Sub9 | L | PHG |
| 17 | Sub10 | L | HPC |
| 18 | Sub10 | L | HPC |
| 19 | Sub1 | R | OFC (4B) |
| 20 | Sub3 | R | OFC (4B, 5B) |
| 21 | Sub4 | R | OFC (4B, 5A) |
| 22 | Sub5 | L | OFC (5B) |
| 23 | Sub6 | L | OFC (3B) |
| 24 | Sub7 | L | OFC (4A) |
| 25 | Sub8 | L | OFC (5B) |
| 26 | Sub9 | L | OFC (5A) |
| 27 | Sub10 | R | OFC (2B) |

